# Sub-2 Å Resolution Structure Determination Using Single-Particle Cryo-EM at 200 keV

**DOI:** 10.1101/855643

**Authors:** Mengyu Wu, Gabriel C. Lander, Mark A. Herzik

## Abstract

Although the advent of direct electron detectors (DEDs) and software developments have enabled the routine use of single-particle cryogenic electron microscopy (cryo-EM) for structure determination of well-behaved specimens to high-resolution, there nonetheless remains a discrepancy between the resolutions attained for biological specimens and the information limits of modern transmission electron microscopes (TEMs). Instruments operating at 300 kV equipped with DEDs are the current paradigm for high-resolution single-particle cryo-EM, while 200 kV TEMs remain comparatively underutilized for purposes beyond sample screening. Here, we expand upon our prior work and demonstrate that one such 200 kV microscope, the Talos Arctica, equipped with a K2 DED is capable of determining structures of macromolecules to as high as ∼1.7 Å resolution. At this resolution, ordered water molecules are readily assigned and holes in aromatic residues can be clearly distinguished in the reconstructions. This work emphasizes the utility of 200 keV for high-resolution single-particle cryo-EM and applications such as structure-based drug design.

---

High-resolution three-dimensional (3D) structure determination of vitrified biological macromolecules by cryogenic electron microscopy (cryo-EM) single-particle analysis (SPA) has expanded the utility of the structural biology field, enabling detailed visualization of targets that were previously intractable using other structural techniques^1-4^. Due to significant advancements in instrumentation, data collection software, and data processing algorithms, obtaining high-resolution reconstructions (i.e., 3 Å resolution or better) has become increasingly routine using SPA, and consequently has garnered substantial interest in the achievable resolution limits of this technique. Indeed, though this resolution regime represents an impressive feat, there remains a considerable discrepancy between what has been presently achieved for structural studies of biological macromolecules (e.g., adeno-associated virus^5^, beta-galactosidase^6,7^, glutamate dehydrogenase^8^, apoferritin^9-11^) versus theoretically attainable resolutions possible with a transmission electron microscope (TEM) (e.g., the Abbe resolution limit given the wavelength of the electron beam). Part of this discrepancy is due to aberrations and distortions inherent to the optics of the instrument, which establish the practical resolution limit in EM^12^. However, ongoing efforts to optimize the integration of aberration correctors with 300 kV TEMs have enabled imaging of materials to resolutions better than 1.0 Å ^13,14^. The remaining gap might be partially abridged through continued technical developments, such as improvements in detector performance or computational correction of aberrations^9,15-18^ but several additional factors currently prevent atomic-resolution cryo-EM SPA of biological specimens. These are largely sample-dependent, and include conformational and/or compositional heterogeneity^19^ and the adoption of preferred orientation, as well as specimen denaturation at the hydrophobic air-water interface^20-23^. Furthermore, high-resolution information of the specimen substantially deteriorates during imaging due to accumulation of beam-induced radiation damage^24^. Advancements in specimen preparation to minimize air-water interface interactions (e.g., Spotiton^25^, graphene or graphene oxide support films^23,26,27^, streptavidin monolayers^28^) and image processing strategies to account for conformational dynamics (e.g., focused classification, multi-body refinement^29^, manifold embedding^30^), are promising avenues that aid in mitigating these oft-encountered issues.

Recent investigations of the resolution limit of cryo-EM SPA have relied upon the use of conformationally homogeneous test specimens that maintain structural integrity through the specimen preparation process. From these efforts, several groups have reported using a TEM operating at 300 kV and equipped with a direct electron detector (DED) to reconstruct biological specimens to better than 2 Å resolution (EMD-0144, EMD-0153, EMD-7770, EMD-8194, EMD-9012, EMD-9599, EMD-9890, EMD-10101, EMD-20026). Several structures of the iron storage protein ferritin in the apo state (apoferritin)^9-11^ have been released with reported resolutions as high as ∼1.5 Å (EMD-9865), attesting to the quality of the test specimen as well as the imaging instrumentation. Indeed, a 300 kV TEM paired with a DED is considered the “industry standard” for high-resolution cryo-EM due to the smaller inelastic scattering cross-section, smaller defocus spread, minimized effects of specimen charging, and a flatter Ewald sphere associated with shorter electron wavelengths^31^. However, the benefits conferred by high-end 300 kV instruments are tied with the high cost of purchase, installation, and maintenance, which are often prohibitive to many institutions or those establishing new cryo-EM facilities^32^. It is therefore prudent to also investigate the resolution capabilities of comparatively more accessible lower-energy TEMs^33,34^, which have not been as extensively characterized for cryo-EM SPA. We have previously shown that targets of varying sizes and symmetries can be determined to high-resolution using a Talos Arctica (operating at 200 kV) equipped with a K2 Summit DED^35,36^. Another group has recently shown that the Thermo Fisher Glacios, which utilizes the same TEM optics as the Arctica, paired with the Falcon 3 DED can be used to determine structures to better than 3 Å resolution^37^. We expand upon our prior work using ∼150 kDa rabbit muscle aldolase by further optimizing our imaging strategies and processing methodologies, including computationally estimating and correcting for higher-order optical aberrations, resulting in an improved ∼2.13 Å reconstruction of this small complex (**Fig. 1**). Further, we demonstrate using mouse heavy chain apoferritin that ∼1.75 Å resolution can be achieved using our instrumentation (**Fig. 2**). At these resolutions, water molecules, coordinated ions, defined backbone features, and holes in aromatic and proline residues can be clearly distinguished in the reconstructions (**Figs. 1-3**). To our knowledge, these are the highest resolutions that have been achieved by cryo-EM SPA using a 200 kV TEM, effectively demonstrating the capacity of lower-energy instruments to break the 2 Å resolution barrier for symmetric, well-behaved specimens.

**Figure 1.**
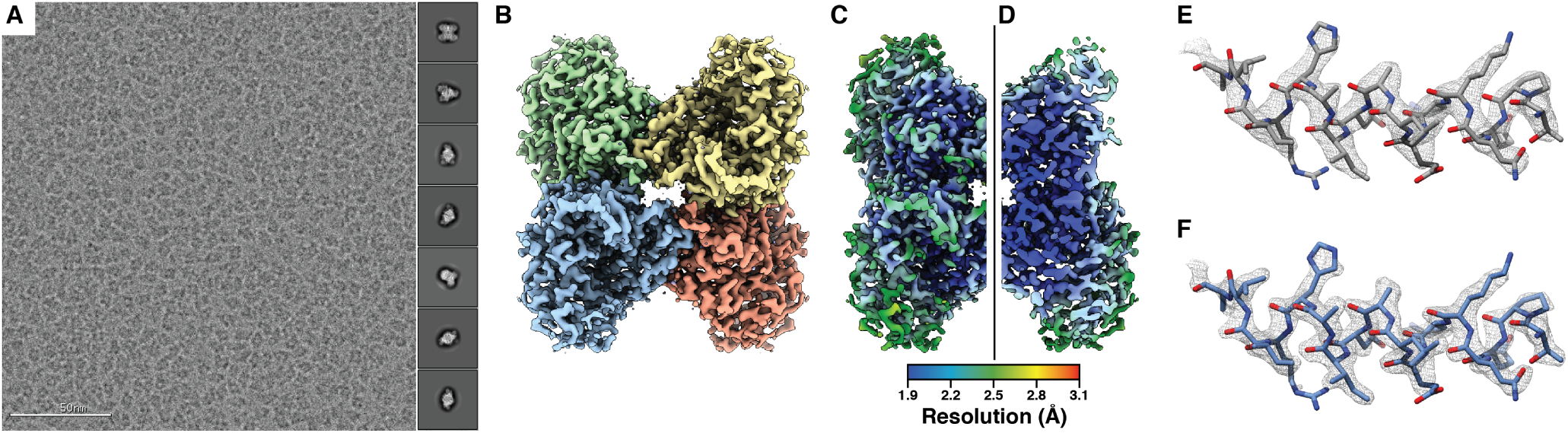
∼2.13 Å resolution cryo-EM reconstruction of 150 kDa rabbit muscle aldolase. **(A)** Representative aligned and dose-weighted micrograph (imaged at ∼1.2 *µ*m underfocus) of aldolase in vitreous ice. Representative reference-free 2D class averages are shown in the right-side inset. **(B)**. Final aldolase EM density colored by subunit. **(C)**. Final aldolase EM density colored by local resolution shown in full or **(D)** sliced in half. **(E)**. and **(F)**. An α-helix comprising residues 8-24 shown in stick representation with EM density (gray mesh) from **(E)** the ∼2.60 Å resolution structure of aldolase36 (EMD-8743) or the **(F)** ∼2.1 Å resolution reconstruction presented here, zoned within 2 Å.

**Figure 2.**
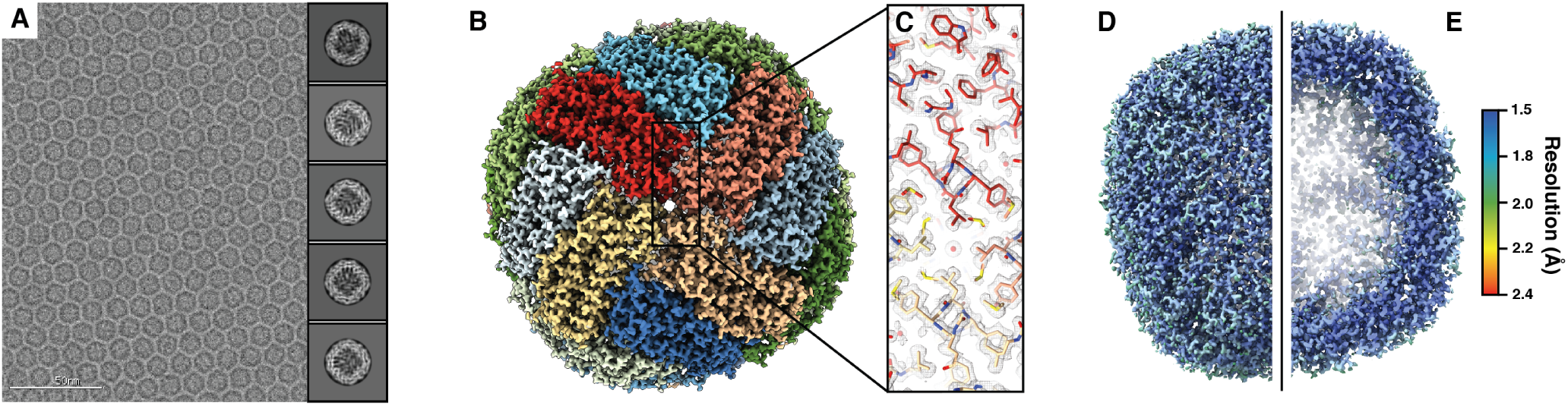
∼1.75 Å resolution cryo-EM reconstruction of mouse heavy chain apoferritin. **(A)**. Representative aligned and dose-weighted micrograph (imaged at ∼1.2 *µ*m underfocus) of apoferritin in vitreous ice. Representative reference-free 2D class averages are shown in the right-side inset. **(B)**. Final apoferritin EM density colored by subunit. **(C)**. Zoomed in region of the final apoferritin EM density (gray mesh). Residues are shown in stick representation and colored by subunit according to **(B). (D)**. Final apoferritin EM density colored by local resolution shown in full or **(E)** sliced in half.

## Results

### Imaging rabbit muscle aldolase and initial reconstruction

We previously determined the ∼2.6 Å resolution structure of rabbit muscle aldolase using a base-model (i.e. excluding imaging accessories such as a phase plate or energy filter) Thermo Fisher Scientific Talos Arctica equipped with a Gatan K2 Summit DED, demonstrating that high-resolution reconstructions are also attainable using a two-condenser lens TEM operating at 200 kV^36^. In our prior study, aldolase was imaged at 45,000x nominal magnification in super-resolution mode and micrographs were Fourier-binned 2 × 2 during motion correction (0.91 Å/physical pixel). Prior detective quantum efficiency (DQE) measurements of a K2 DED at 200 keV showed a fall below 0.5 at half Nyquist frequency in both counting and super-resolution imaging modes^38^. Intriguingly, these data also demonstrated that the DQE below half Nyquist frequency is higher in counting mode than in super-resolution mode. Given these results, we attempted to further improve the resolution of aldolase by conducting additional studies in counting mode at a higher magnification (73,000x nominal magnification, corresponding to 0.56 Å/physical pixel) to utilize the higher DQE within our resolution regime. TEM column alignments were performed as described previously^35^, with modifications to further maximize parallel illumination using a longer diffraction mode camera length (see **Methods**). Iterative 3D classification and refinement steps were performed with D2 symmetry imposed throughout, followed by Bayesian particle polishing and refinement of defocus UV and global astigmatism using RELION 3.0^9^, yielding a ∼2.3 Å reconstruction (see **Methods** and **SI Fig. 1**).

### Imaging mouse apoferritin and initial reconstruction

To further complement our studies as well as those performed using 300 kV TEMs, we sought to investigate the resolution limits of our instrumentation using a more conformationally homogeneous and symmetric test specimen. We thereby selected mouse heavy-chain apoferritin, a ∼505 kilodalton (kDa) complex with octahedral symmetry. Movies of vitrified apoferritin were collected at 73,000x magnification using a Talos Arctica TEM equipped with a K2 Summit DED (see **Methods**). In anticipation of obtaining a nominal resolution approaching Nyquist frequency due to the exceptional structural stability of apoferritin, we elected to collect images using super-resolution mode to better preserve the high-resolution information beyond Nyquist. The majority of collected images contained a single layer of apoferritin particles in a pseudo-lattice arrangement, indicating the molecules were embedded in a thin layer of vitreous ice (**Fig. 2A**). Appion^39^ and RELION 3.0^9^ were used for image processing. Reference-free two-dimensional (2D) classification and iterative rounds of 3D refinement and CTF refinement (per-particle defocus, global astigmatism, beam tilt) with octahedral symmetry applied yielded a ∼2.2 Å reconstruction, and Bayesian particle polishing improved the resolution to ∼2.0 Å. Successive attempts at 3D classification and CTF refinement did not improve the nominal resolution nor the visual quality of the reconstruction (see **Methods** and (**SI Fig. 3**).

**Figure 3.**
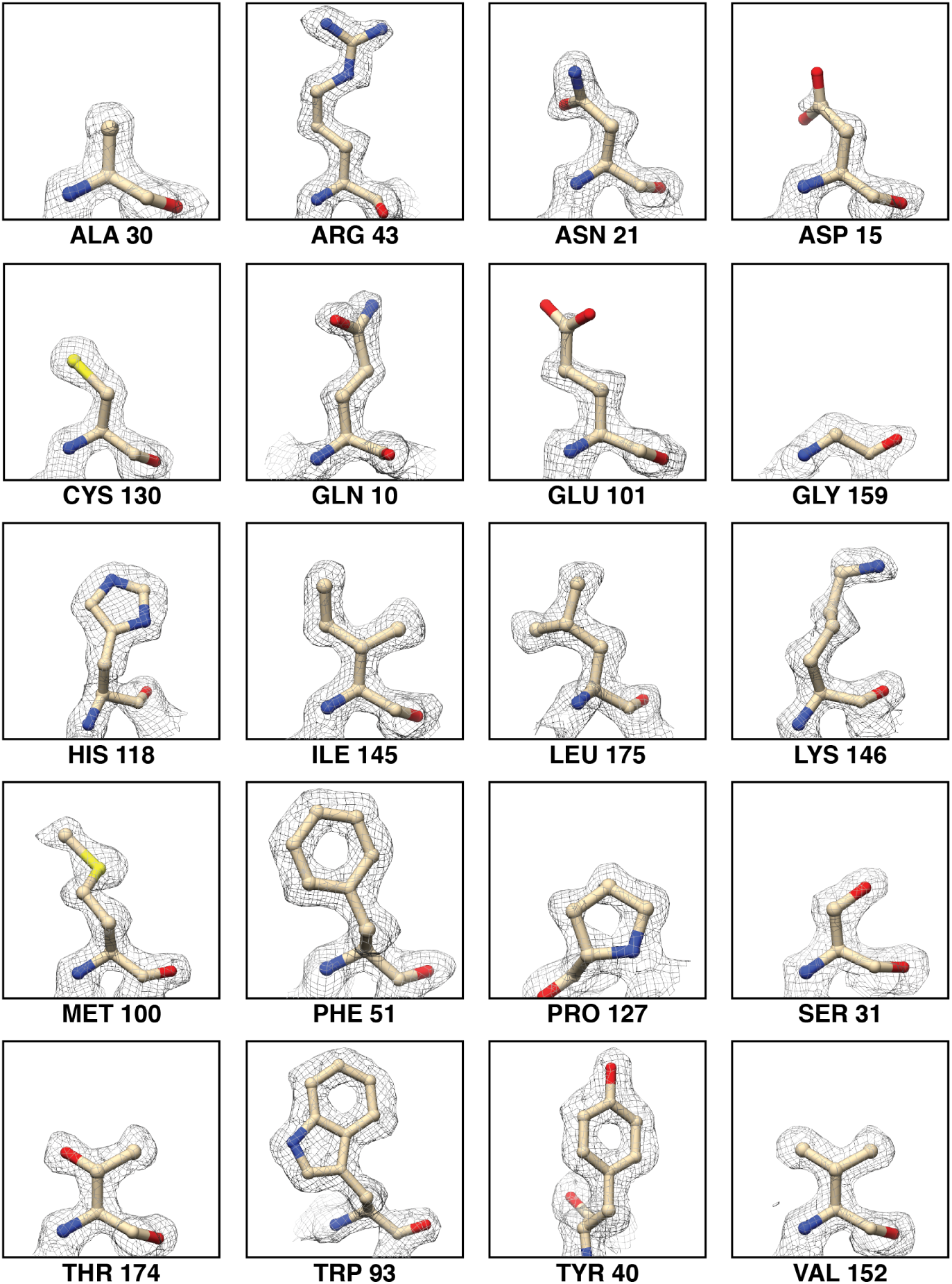
Apoferritin EM map quality for each of the standard 20 amino acids. The EM density zoned 2 Å around the residue atoms (shown in stick representation) is shown in gray mesh. All amino acids are shown at the same map contour level. The corresponding residue number is shown below each panel.

Our inability to improve the resolutions of the reconstructions beyond 2 Å using established reconstruction methodologies compelled us to investigate the extent to which inaccuracies in the magnification/pixel size and/or the typically fixed values supplied by instrumentation vendors that are used for CTF estimation, such as spherical aberration of the objective lens, might be impacting our ability to achieve resolutions better than 2 Å.

### Magnification refinement

The nominal voxel size value used in this study (0.56 Å/pixel) was extrapolated from manually measured distances between diffraction spots of a single gold crystal at a lower magnification, and as such is only expected to be accurate to ±5-15%^40^. Further, these values can be influenced by variability in TEM optics as well as radiation-induced distortions of the biological specimen^40^. Given that vitrified specimens prepared for cryo-EM retain inter-atomic distances that are more consistent with those measured in room temperature X-ray diffraction studies^40-42^, we investigated alternative methods to identify a more accurate magnification. To accomplish this, the X-ray crystal structures of the asymmetric units of apoferritin (PDB ID: 3WNW) or aldolase (PDB ID: 5VY5, that was subjected to a short molecular dynamics simulation, see Methods) were rigid-body refined into respective EM reconstructions of voxel sizes ranging from 0.555 Å/pixel to 0.575 Å/pixel using the Phenix.real space refinement package^43^. Fitting polynomial curves to the plots of local model-map cross-correlation (CC) against voxel size indicated a clear maximum correlation at 0.561 Å/pixel for apoferritin and 0.563 Å/pixel for aldolase (**SI Fig. 5**). The average of these values (0.562 Å/pixel) was taken to be the refined voxel size for both data sets.

### Spherical aberration refinement

The spherical aberration (C_s_) value directly contributes to phase error during imaging and becomes an important component of the contrast transfer function (CTF) at high spatial frequencies. The C_s_ value is supplied by the microscope manufacturers and is typically not refined during image processing, though it is conceivable that there exists some variability across lenses in different systems. We therefore set out to investigate if changing the C_s_ value of 2.70 mm ascribed to Thermo Fisher Scientific TEMs such as the Titan Krios, Talos Arctica, and Glacios would impact our attainable resolution. Given that available protocols used to measure the C_s_ of our Arctica would yield values with error ranges of ±10% due to errors in measuring focus, wavelength, and magnification^44^, we instead opted to refine a subset of “polished” apoferritin particles using C_s_ values ranging from 2.60 mm to 2.95 mm (see **Methods**). The highest resolution reconstruction (∼2.11 Å) was obtained when “correcting” with C_s_=2.80 mm (**SI Fig. 4**). When we used these optimized instrument values (0.562 Å/pixel, C_s_=2.80 mm) to reprocess the entire dataset of particles, we were able to further improve the resolution of apoferritin to a global resolution of ∼1.8 Å (**SI Fig. 3**). However, performing the aforementioned C_s_ refinement procedure for aldolase using C_s_ values other than 2.70 mm did not yield significant improvements in resolution or quality of the resulting reconstructions (data not shown). Because the phase shifts arising from C_s_ increase as a fourth power of the spatial frequency, we speculate that the resolution of aldolase (∼2.3 Å) was not sufficiently high enough to be noticeably influenced by an inaccuracy in the C_s_.

### Quantification and correction of higher-order aberrations

Given that changing the C_s_ parameter value improved the resolution for apoferritin, but not for aldolase, we could not eliminate the possibility that by manipulating this parameter we were likely also accommodating for unexplored higher-order optical aberrations that were present in the dataset. Unlike the Titan Krios, two-condenser systems such as the Arctica or Glacios do not contain multipole correctors to physically compensate for these aberrations (e.g., a hexapole corrector, which can be used to minimize three-fold astigmatism). We computationally characterized the higher-order aberrations present in our data using RELION 3.1, which models the antisymmetrical and symmetrical components using the first six anti-symmetrical Zernike polynomials (radial orders 1 and 3) and the first nine symmetrical polynomials (radial orders 0, 2, and 4)^15^.

Our aldolase data exhibited both anti-symmetrical and symmetrical aberrations, as evidenced by the resulting per-Fourier-pixel average phase-error plots (**SI Fig. 2**). The anti-symmetrical aberrations corresponded to the presence of beam tilt and axial coma (but minimal three-fold/trefoil astigmatism). The symmetrical aberration plots indicated strong four-fold/tetrafoil astigmatism in both data sets, particularly for the second subset. The symmetrical aberrations were estimated while keeping C_s_ fixed at 2.70 mm. To further refine the C_s_ (also a 4th order symmetrical aberration), we allowed per-micrograph fitting of C_s_ values using input values of 2.70 mm or 2.80 mm. The C_s_ estimates fit from both inputs converged on the same mean value of 2.7 mm ± 0.02 mm, in agreement with the results of our spherical aberration refinement and the default specifications for this parameter. The aldolase data also contained weak magnification anisotropy (**SI Fig. 2**). Correcting for these aberrations further improved the resolution of the aldolase reconstruction to ∼2.13 Å (**Fig. 1B, SI Fig. 1**, and **Table 1**). Local resolution estimation of the resulting reconstruction revealed the majority of the core of molecule was resolved to ∼1.90 Å (Fig. 1, C and D). Comparison of this reconstruction to EMD-8743 revealed better-defined backbone and side-chain density overall (Fig. 1, E and F).

The anti-symmetrical aberration plot of the apoferritin data also revealed some beam tilt and axial coma, but again no significant trefoil astigmatism. The symmetrical aberrations corresponded to pronounced tetrafoil astigmatism (**SI Fig. 3**). Interestingly, when we performed per-micrograph fitting of C_s_ as was done for aldolase, the estimates did not deviate much from the input value (i.e. the 2.70 mm and 2.80 mm inputs returned mean values around 2.7 mm and 2.8 mm, respectively). However, the resulting estimated symmetrical Zernike polynomials changed substantially between these refinements, indicating that other terms could be absorbing errors in the C_s_ (and vice-versa). Specifically, as the global average of the C_s_ is absorbed by the 0th and 4th terms of the symmetrical polynomials, changes in these values could produce the same “effective” C_s_ if the input value is incorrect. Given that these polynomial terms were smaller for the 2.80 mm refinements than for the 2.70 mm refinements (1.09 and 4.03, respectively), we reason that the actual C_s_ of our microscope likely differs from the factory-specified value of 2.70 mm (**SI Fig. 3**). However, the resulting aberration-corrected apoferritin reconstructions using C_s_ values refined from 2.70 mm or 2.80 mm were very similar both in terms of resolution (∼1.78 Å and 1.75 Å, respectively) and quality, suggesting that tetrafoil and errors in C_s_ may have been compensated for by other symmetrical (possibly even higher-order) aberrations.

Inspection of the ∼1.75 Å reconstruction revealed defined backbone carbonyl groups, clear density for ordered water molecules and coordinated zinc atoms, as well as distinct side-chain structural details such as holes in aromatic rings and notably greater density associated with sulfur atoms, all of which attest to the quality of the map (Fig. 2, B and C; **Fig. 3**). Local resolution estimation of the reconstruction revealed a fairly uniform resolution distribution, with the core of the ferritin shell resolved to ∼1.6 Å (Fig. 2, D and E). Taken together, these results demonstrate that higher-order aberrations that contribute to phase errors and distortions of high-frequency information in our data can now be computationally modeled and corrected for in the absence of multipole aberration correctors, and advances in software will streamline this process. Importantly, CTF parameter values that were previously believed to be invariant for a given type of microscope, such as C_s_, may indeed vary from factory specifications, and refinement of these values concurrent with estimation of higher order aberrations could yield improvements in nominal resolution.

### Acceleration voltage refinement

Given that the CTF depends on both the spherical aberration coefficient and the electron wavelength (as defined by the accelerating voltage), we also explored whether the acceleration voltage of the microscope might differ from the factory-specified value of 200 kV for our apoferritin data set. We used the same refinement procedure outlined for estimating the C_s_ value to determine the acceleration voltage that produces the highest resolution structure from our data. Briefly, we refined a subset of “polished” apoferritin particles using accelerating voltage values ranging from 194 to 206 kV in 2 kV increments (see Methods). Particles that utilized a C_s_=2.70 mm returned a value of 196 kV, while particles that utilized a C_s_=2.80 mm returned a value of 200 kV and refined to a slightly improved nominal resolution (∼2.11 Å) than that of a reconstruction obtained from C_s_=2.70 mm particles (∼2.14 Å) (**SI Fig. 4**). Although we cannot eliminate the possibility that the voltage may have varied during data collection, these refinements indicate that our acceleration voltage did not significantly deviate from 200 kV.

## Concluding remarks

Despite the ubiquity of single particle cryo-EM, there is a lingering misconception that high-resolution structure determination of biological targets can only be achieved with 300 kV three-condenser lens instruments such as the Titan Krios, with 200 kV instrumentation primarily relegated to sample screening. Given the costs associated with establishing and maintaining 300 kV systems, it is not only prohibitive but also impractical to rely solely on 300 kV TEMs to meet the demands of the structural biology community. Our work demonstrates that a two-condenser lens TEM operating at 200 kV equipped with a DED can produce SPA cryo-EM reconstructions of ideal biological specimens to resolutions better than 2 Å. Given that the vast majority of SPA cryo-EM structures deposited in the EMDB are not resolved beyond 2 Å resolution, we posit that issues associated with specimen preparation and/or sample heterogeneity, rather than microscope optics or electron detection, are the greatest limiting factor to the attainable resolution of SPA reconstructions.

A notable drawback to 200 kV instruments is the lack of aberration correctors, which contributes to the lower cost of the instruments but consequently introduces the likelihood of phase incoherence at high spatial frequencies. Our studies show that trefoil and tetrafoil aberrations can be computationally estimated and modeled using RELION 3.1^15^ to obtain maps containing clear density for coordinated ligands, ordered water molecules, and, particularly in the case of the apoferritin reconstruction, distinct holes in aromatic and proline residues. The capacity of a 200 kV instrument to distinguish these features holds great promise for its applications in high-resolution structure determination as well as structure-based drug design. Curiously, the extent of these aberrations (which would be expected to remain consistent for a given instrument) seemed to vary between imaging sessions as well as between specimens (**SI Figs 2** and **3**). These inconsistencies warrant future investigations into the cause of the aberrations observed on our Arctica. We also cannot exclude the contributions of other main axial aberrations within our instrument such as other higher-order aberrations as well as chromatic aberration (C_c_), the effects of which are stronger at lower accelerating voltages^45^. As such, in lieu of an experimental means to mitigate these, it is necessary to utilize *in silico* methods for estimating and correcting for such deviations as demonstrated here and elsewhere^15,16^. In addition, we speculate that refinement of CTF parameter values that had previously been deemed invariant from the factory specifications will also lead to improvements in cryo-EM reconstructions and better characterization of TEMs in operation. Together, we anticipate that the advent of such computational tools and refinement procedures will greatly benefit the EM community by not only informing on instrument aberrations, but also by expanding the resolving capabilities of TEMs for cryo-EM SPA of biological complexes.

## Materials and Methods

### Sample preparation

#### Rabbit muscle aldolase

Lyophilized rabbit muscle aldolase (Sigma-Aldrich) was solubilized in 20 mM HEPES pH 7.5, 50 mM NaCl to a final concentration of ∼3 mg/ml further purified by size-exclusion chromatography using a Sepharose 6 10/300 (GE Healthcare) column equilibrated in solubilization buffer. Fractions containing the highest purity aldolase, as determined by SDS-PAGE, were pooled and concentrated to ∼1.6 mg/ml.

#### Mouse heavy chain apoferritin

A pET24a vector encoding the heavy chain of mouse apoferritin was received from M. Kikkawa (The University of Tokyo) and, without modification, was transformed into BL21(DE3)pLys *E. coli* chemically-competent cells. Cells were grown at 37° C in LB media until an OD_600nm_ = 0.5 was reached. Protein expression was induced with 1 mM IPTG at 37° C. After 3 hours, cells were pelleted at 4,000× g for 10 min (4 ° C), resuspended in lysis buffer (30 mM HEPES pH 7.5, 300 mM NaCl, 1 mM MgSO_4_) supplemented with 1 mg/ml lysozyme and cOmplete Protease Inhibitor Cocktail (Roche), and subsequently lysed using sonication. Cell debris were pelleted at 20,000×g for 30 min (4° C) and the clarified supernatant was heat-treated at 70° C for 10 minutes to denature endogenous *E. coli* proteins. Denatured proteins were pelleted at 20,000×g for 15 minutes (4° C) and ammonium sulfate was then added to the cleared supernatant to a final concentration of 60% (w/v) followed by gentle stirring on ice for 10 minutes. The precipitant was harvested at 14,000×g for 20 minutes (4° C), gently resuspended in 2 mL of cold phosphate buffered saline (PBS) and subsequently dialyzed against buffer Q1 (30 mM HEPES pH 7.5, 1 mM DTT, 20 mM NaCl). Dialyzed protein was diluted two-fold in buffer Q1 and loaded onto a HiTrap Q HP anion exchange chromatography column (GE Healthcare), equilibrated in buffer Q1, at 1 mL/min. The column was washed with 4 column volumes of buffer Q1 and protein was eluted using a 0-100% gradient of buffer Q2 (30 mM HEPES pH 7.5, 1 mM DTT, 500 mM NaCl) applied over 3 column volumes. Apoferritin eluted between 150-200 mM NaCl, as confirmed by SDS-PAGE. Samples corresponding to pure apoferritin were pooled, concentrated to 10-20 mg/mL, and loaded onto a Superdex 200 Increase 10/300 (GE Healthcare) size exclusion chromatography column equilibrated with 30 mM HEPES pH 7.5, 150 mM NaCl, 1 mM DTT. Peak fractions corresponding to the highest purity apoferritin were pooled and concentrated to 4-5 mg/mL. For long-term storage, trehalose (5% (v/v) final) was added to concentrated apoferritin prior to flash-freezing in liquid nitrogen.

### Cryo-EM grid preparation

3 *µ*L of purified aldolase (1.6 mg/mL) or apoferritin (5 mg/mL) was dispensed on UltrAuFoil R1.2/1.3 300-mesh grids (Quantifoil Micro Tools GmbH) that had been freshly plasma cleaned for six seconds at 15Watts (75% nitrogen/25% oxygen atmosphere) using a Solarus plasma cleaner (Gatan, Inc.). Grids were manually blotted for four to five seconds using Whatman No. 1 filter paper and immediately plunge-frozen into liquid ethane cooled by liquid nitrogen using a custom-built manual plunger located in a cold room (^≥^95% relative humidity, 4° C).

### Cryo-EM data acquisition, image processing, and refinement

Microscope alignments were performed on a crossed-lines grating replica calibration grid using previously described methodologies, including determining parallel illumination using a long diffraction camera length^35,46^. Movies of frozen-hydrated aldolase or apoferritin were collected using a Talos Arctica transmission electron microscope (Thermo Fisher Scientific) operating at 200 keV and equipped with a K2 Summit direct electron detector (Gatan, Inc.). All cryo-EM data were acquired using the Leginon automated data collection software^47^ and pre-processed in real-time using the Appion package^39^. Movies of aldolase and apoferritin were collected in counting mode (0.56 Å/pixel) and super-resolution mode (0.28 Å/super-resolution pixel), respectively, at a nominal magnification of 73,000x over a defocus range of −0.3 *µ*m to −1.8 *µ*m. Aldolase movies were obtained over two imaging sessions, approximately 1 month apart, both using an exposure rate of 1.92 e^-^/pixel/s for a total of 11 seconds (250 ms/frame, 44 frames), resulting in a total exposure of ∼67 e^-^/Å^2^ (1.52 e^-^/Å^2^/frame). Apoferritin movies were obtained using an exposure rate of ∼2 e^-^/pixel/s for a total of 9 seconds (100 ms/frame, 90 frames), resulting in a total exposure of ∼58 e^-^/Å^2^ (0.64 e^-^/Å^2^/frame). The apoferritin frames were Fourier-binned 2 × 2 (0.56 Å/pixel) prior to motion correction using the MotionCor2 frame alignment program^48^ implemented within RELION 3.0v^29^. Frame alignment without dose-weighting was performed on 4 × 4 tiled frames with a B-factor of 250 and a running average of 2 frames for both aldolase and apoferritin images. Aligned frame stacks were also saved to be later used for Bayesian Polishing^9^. Un-weighted, summed images were used for local CTF estimation using equiphase averaging (512-pixel local box size, 0.10 amplitude contrast, 30 Å minimum resolution, 3 Å maximum resolution)^49^. Aligned images with a CTF maximum resolution worse than 4 Å were excluded from further processing.

#### Aldolase

For aldolase, EMD-8743 was low-pass filtered to 20 Å and used to generate 2D templates for automated template-based particle picking of both datasets using RELION 3.0v2. A total of 1,801,738 particles picks were extracted from 3,534 micrographs collected across both sessions, binned 4 × 4 (2.24 Å/pixel, 128 pixel box size) and subjected to reference-free 2D classification (200 classes, tau fudge=2, 120 Å mask diameter). Particles corresponding to 2D class averages containing strong secondary structural details were isolated (982,098 particles) and 3D auto-refined with D2 symmetry using EMD-8743 low-pass filtered to 30 Å as an initial model. The refined coordinates were used to re-center and re-extract particles binned 2 × 2 (1.12 Å/pixel, 256-pixel box size). These particles were refined again using a scaled version of the previously refined map, followed by 3D classification (4 classes, tau fudge=2, E-step limit=7 Å) using a soft mask (5-pixel extension, 5-pixel soft cosine edge). A total of 394,294 particles were selected from the 3D classes and subjected to per-particle CTF refinement (defocus and global astigmatism) and beam tilt estimation. The particles were then 3D auto-refined, re-centered, and re-extracted without binning (0.56 Å/pixel, 256-pixel box size). Particle motion trajectories and radiation damage were estimated using RELION 3.0v2 Bayesian particle polishing. Shiny particles were re-extracted without binning (512-pixel box size) and 3D auto-refined to ∼2.4 Å resolution. An additional iteration of CTF refinement (defocus and global astigmatism) was performed and beam tilt was estimated for the two datasets separately (for the first dataset, x=0.23 mrad, y=-0.08 mrad; for the second dataset, x=-0.17 mrad, y=0.14 mrad). 3D auto-refinement yielded a ∼2.3 Å resolution reconstruction (∼2.9 Å with C1 symmetry). Estimation of magnification anisotropy and higher-order optical aberrations (anti-symmetrical and symmetrical components) was performed using RELION 3.1^15^. Per-micrograph refinement of C_s_ was also performed (2.7 ^±^ 0.02 mm). Correcting for these aberrations yielded a final ∼2.1 Å resolution reconstruction according to gold-standard FSC^50^. Local resolution estimation was calculated using the blocres function in BSOFT^51^.

#### Apoferritin

For apoferritin, RELION 3.0v2 was used to manually pick particles from the first 50 micrographs, yielding 1,102 picks, that were then extracted binned 2 × 2 (1.12 Å/pixel, 192-pixel box size) and subjected to reference-free 2D classification. Nearly all particles converged to a single 2D class which was used for automated template-based particle picking against the entire set of aligned images using RELION. A total of 405,106 picks were extracted from 1759 micrographs, binned 2 × 2 (1.12 Å/pixel, 192-pixel box size), and subjected to reference-free 2D classification (50 classes, tau fudge=2). Particles corresponding to 2D class averages containing strong secondary structural details were isolated (158,138 particles) and the remaining particles were subjected to an additional round of 2D classification (25 classes, tau fudge=2, ignoring information until the first CTF peak), from which 135,116 particles were isolated. This procedure was repeated once more (yielding 30,217 particles) and all selected particles were combined to yield 323,471 particles in total. These particles were subjected to RELION 3D auto-refinement with O symmetry using an initial model generated from EMD-9599 low-pass filtered to 8 Å to provide enough information to assign the orientation of the alpha helices^52^. The refined coordinates were used for subsequent re-centering and re-extraction of unbinned particles (0.56 Å/pixel, 384-pixel box size), which were refined using a scaled version of the previous map to yield a ∼2.4 Å reconstruction (gold-standard FSC at 0.143 cutoff).

Refinement of per-particle defocus, beam tilt, and global astigmatism were performed using RELION 3.0v2, followed by 3D auto-refinement yielding a ∼2.2 Å reconstruction. The particles were then subjected to RELION Bayesian particle polishing, which further improved the resolution of the reconstruction to ∼2.1 Å. Repeating per-particle defocus, beam tilt, and per-particle astigmatism refinement yielded a ∼2.0 Å reconstruction. The particles were then subjected to no-alignment 3D classification (4 classes, tau fudge=12) using a soft mask (5-pixel extension, 10-pixel soft cosine edge), which yielded a single well-resolved class comprising 323,362 particles. It was previously observed that eliminating particles imaged at higher underfocus yielded modest improvements in nominal resolution and map quality^35,36^. From this reasoning, we selected only those images collected between 200 - 12000 nm underfocus (241,878 particles) for 3D auto-refinement; however, no improvement in resolution was observed. Further attempts at 3D classification, particle sorting (e.g., defocus values, Z-scores, rlnNrSignificantSamples, etc.), or modifications to 3D auto-refinement parameters did not improve the nominal FSC-reported resolution beyond ∼2 Å. To explore the possibility that errors in pixel size or spherical aberration were limiting our ability to attain a higher resolution reconstruction, we refined these values using the methodologies described in the subsequent sections.

In accordance with the results of the C_s_ refinement tests, the particles were refined with C_s_=2.80 mm, yielding a ∼1.89 Å reconstruction. After repeating per-particle defocus and astigmatism refinement with beam tilt estimation (x=0.15 mrad, y=-0.14 mrad), the resolution of the resulting reconstruction improved to ∼1.84 Å. Estimation of higher-order optical aberrations (anti-symmetrical and symmetrical components) and per-micrograph C_s_ refinement were performed using RELION 3.1^15^ for particle stacks with C_s_=2.80 mm or 2.70 mm (yielding values of 2.8 ^±^ 0.01 mm and 2.7 ^±^ 0.006 mm, respectively). Correcting for these aberrations using refined C_s_=∼2.8 mm or ∼2.7 mm yielded reconstructions at ∼1.75 Å and ∼1.78 Å resolutions, respectively, according to gold-standard FSC^50^. Local resolution estimation of the 1.75 Å was calculated using the blocres function in BSOFT^51^.

### Pixel size refinement

X-ray crystal structures of apoferritin (3WNW) and aldolase (5VY5) asymmetric units were rigid-body refined into the EM reconstructions determined using voxel sizes between 0.555 Å/pixel and 0.575 Å/pixel using the Phenix.real space refinement package^43^. For aldolase, 5VY5 was subjected to a short molecular dynamics simulation to minimize bias in the starting model from compaction upon freezing of the specimen for X-ray diffraction studies. Briefly, PDB2QR^53^ was used for the initial assignment of charges of aldolase and the CHARMM-GUI^54,55^ was used to generate the input files for NAMD^56^. Aldolase was then subjected to a standard equilibration followed by a short 20 ns simulation run using implicit solvent. The outputted snapshots (every 1 ns) were monitored to ensure protein structural integrity was maintained and the PDB file of the final snapshot was used for voxel size refinement. Second-order polynomial curves were fit to the plot of local model-map cross-correlation (CC) against voxel size for each specimen (**SI Fig. 5**).

### Spherical aberration (C_s_) refinement

For apoferritin, particles from the first 50 micrographs (11,902 particles) of the final refined particle stack that yielded the ∼2 Å resolution reconstruction were selected for testing CTF parameters. The RELION star file representing these particles was manually modified to change the C_s_ value from 2.70 mm (factory specifications) to values between 2.60 – 2.95 mm in 0.05 mm increments. The modified star files were then used for defocus refinement as implemented within RELION while keeping the CTF astigmatism and azimuth angle values constant. The star file containing updated defocus estimates were then subjected to 3D auto-refinement with O symmetry enforced using the 2 Å resolution reconstruction low-pass filtered to 8 Å as an initial model. Each reconstruction was postprocessed within RELION, subjected to an additional round of CTF refinement and subsequent 3D auto-refinement. The final FSC-estimated resolution for each reconstruction was then plotted against C_s_ value and a second order polynomial was fitted (**SI Fig. 4**). A minimum was observed at C_s_=2.80 mm. The RELION star file representing all the particles contributing to the ∼2 Å reconstruction was manually modified to change the C_s_ value to 2.80 mm and subjected to further CTF refinement and 3D auto-refinement (see above).

### Acceleration voltage refinement

The same subset of “polished” apoferritin particles previously used for C_s_ refinement were manually modified to change the accelerating voltage from 200 kV to values ranging from 194 – 206 kV in 2 kV increments. The C_s_ value of these subsets were set to 2.70 mm or 2.80 mm. The same refinement procedure was performed as described above. The final FSC-estimated resolution was plotted against accelerating voltage value for each C_s_ value, and a second order polynomial was fitted (**SI Fig. 4**). A minimum was observed at 196 kV and 200 kV for subsets using C_s_=2.70 mm and 2.80 mm, respectively.

## Acknowledgements

We thank J.-C. Ducom at The Scripps Research Institute (TSRI) High Performance Computing for computational support, B. Anderson at the TSRI microscopy facility for microscope support, and J. Zivanov, T. Nakane, and S. Scheres for insightful discussion and early access to RELION 3.1. We are grateful to M. Kikkawa and H. Yanagisawa for providing the mouse apoferritin plasmid and expression/purification protocols. M.W. is supported as a National Science Foundation Graduate Student Research fellow. G.C.L. is supported by a young investigator award from Amgen, and by the NIH (DP2EB020402 and R21AR072910). M.A.H. was partially supported as a Helen Hay Whitney Foundation Postdoctoral Fellow. Computational analyses of EM data were performed using shared instrumentation funded by NIH S10OD021634 and the molecular dynamics simulations were performed using a private cluster provided as part of a startup package to M.A.H.

## Data availability

The atomic coordinates for the aldolase and apoferritin structures have been deposited in the Protein Data Bank (PDB) under accession codes 6V20 and 6V21, respectively. The corresponding EM density maps (final unsharpened and sharpened maps, half maps, and masks) have been deposited to the Electron Microscopy Data Bank under accessions EMD-21023 and EMD-21024 for aldolase and apoferritin, respectively. Uncorrected movie frames and associated gain correction images for the aldolase and apoferritin datasets in this study have been deposited to the EMPIAR database under accessions 10338 and 10337, respectively.

## Supplementary Materials

**Supplementary Figure 1.**
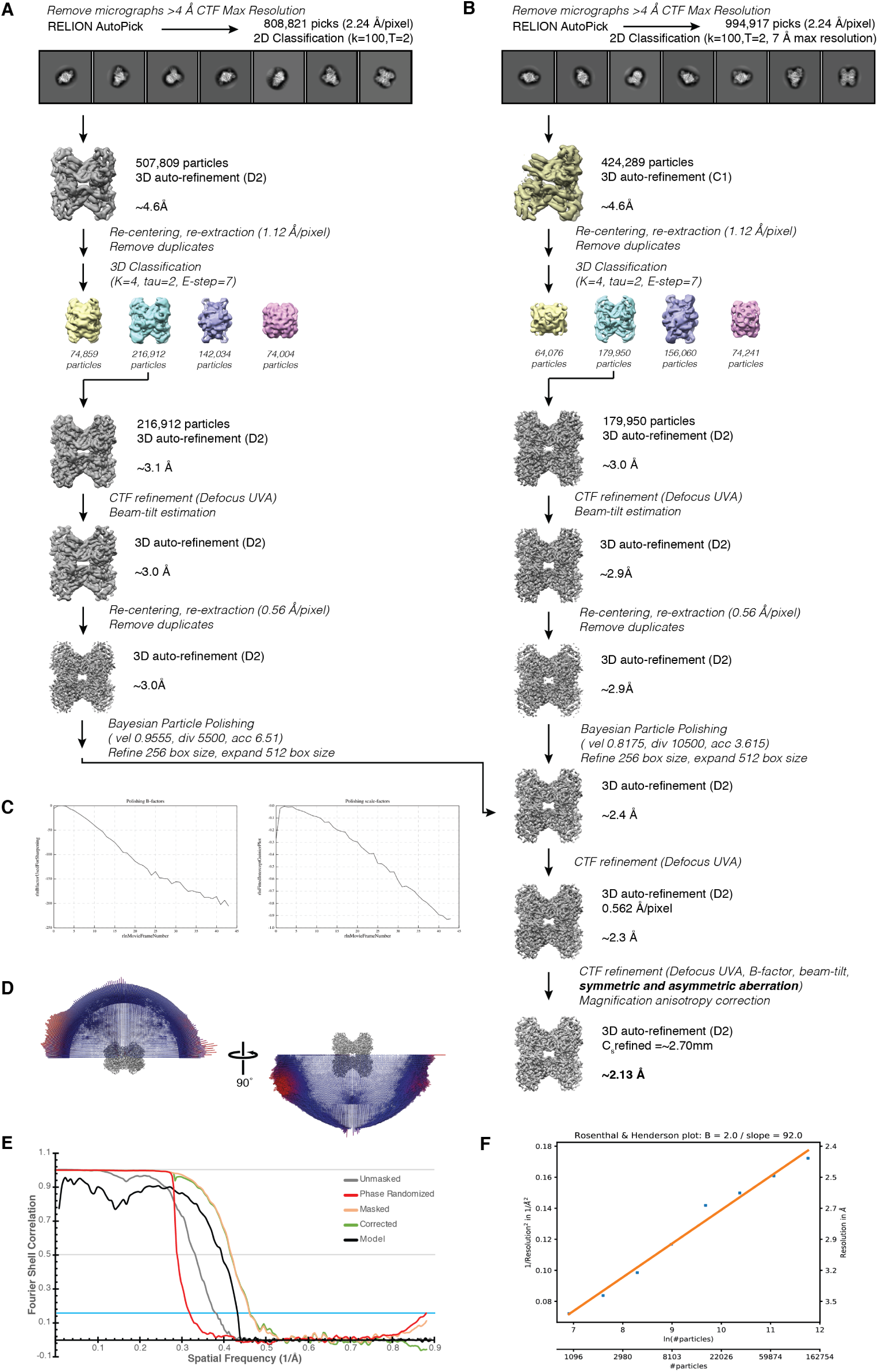
Schematic for aldolase single-particle cryo-EM data processing. ∼ 800K particles (**(A)** and ∼990K particles **(B)** were extracted from aligned, non-dose-weighted micrographs, Fourier-binned 4 × 4 (64 pixel box, 2.24 Å/pixel), and subjected to two independent rounds of reference-free 2D classification using RELION 3.0^1^. Representative class averages are shown. Particles comprising the “best” classes were 3D auto-refined to ∼4.6 Å resolution. Particles were re-centered and re-extracted Fourier-binned 2 × 2 (128 pixel box, 1.12 Å/pixel), and subjected to 3D classification (K=4, tau fudge=2, E-step=7 Å). ∼217K **(A)** and ∼180K **(B)** particles comprising the best class were selected for 3D auto-refinement followed by CTF refinement (defocus UV, whole micrograph astigmatism), including beam-tilt estimation, and another round of 3D auto-refinement. Particles were re-centered and re-extracted (256 pixel box, 0.56 Å/pixel), subjected to 3D auto-refinement, and the outputs were used for Bayesian particle polishing to yield “shiny” particles (512 pixel box, 0.56 Å/pixel). Plots of the calculated B-factors and scale factors from Bayesian particle polishing for dataset **(B)** are shown in **(C)**. The outputted “shiny” particles from both datasets were combined (∼390K particles) and subsequently 3D auto-refined to yield a ∼2.4 Å resolution structure. CTF refinement (defocus UV, whole micrograph astigmatism), and subsequent 3D auto-refinement yielded a ∼2.3 Å resolution structure. An additional round of CTF refinement including higher-order optical aberration correction and Cs refinement yielded a ∼2.1 Å resolution reconstruction. **(D)** Plots showing the Euler angle distribution of the final aldolase EM density. **(E)** Gold-standard Fourier shell correlation (FSC) curve generated from the independent half maps contributing to the ∼2.1 Å resolution aldolase EM density. FSC curve between the final refined atomic model and aldolase EM density is also shown. **(F)** Rosenthal and Henderson plot for particles contributing to the final aldolase reconstruction.

**Supplementary Figure 2.**
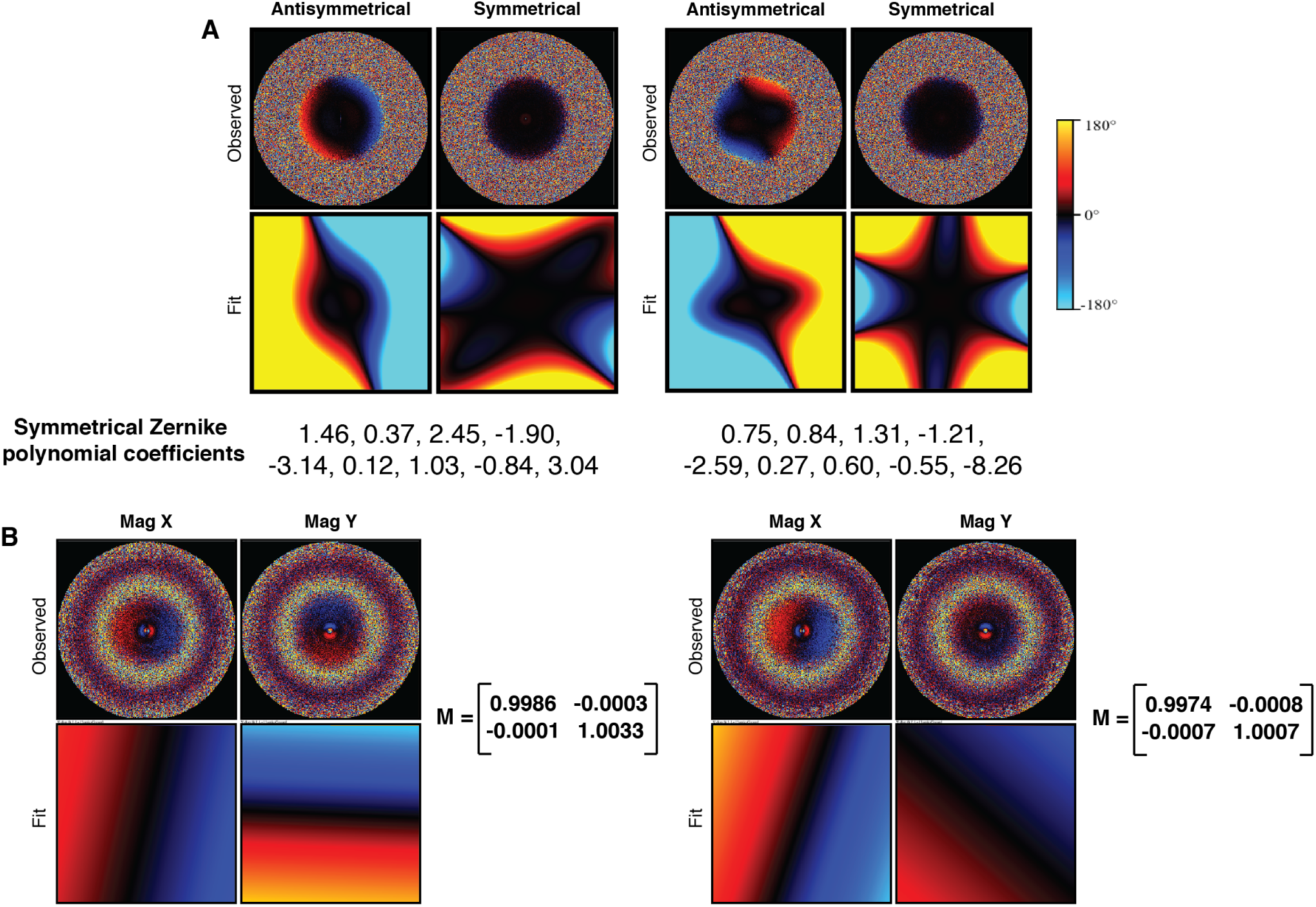
Estimated aberrations and magnification anisotropy for aldolase datasets. **(A)** Anti-symmetrical and symmetrical aberrations estimated for both datasets (left, dataset corresponding to **(SI Fig. 1A)**; right, dataset corresponding to **(SI Fig. 1B)**) using RELION 3.1^2^. Per-pixel phase-angle estimates are depicted in the upper plots, and parametric fits using Zernike polynomials are shown in the lower plots. The estimated symmetrical Zernike polynomial coefficients are shown below each panel. **(B)** Magnification anisotropy estimated for both datasets. Observed and fit per-pixel displacements are depicted in the upper and lower plots, respectively, and the estimated magnification matrix M is included on the right.

**Supplementary Figure 3.**
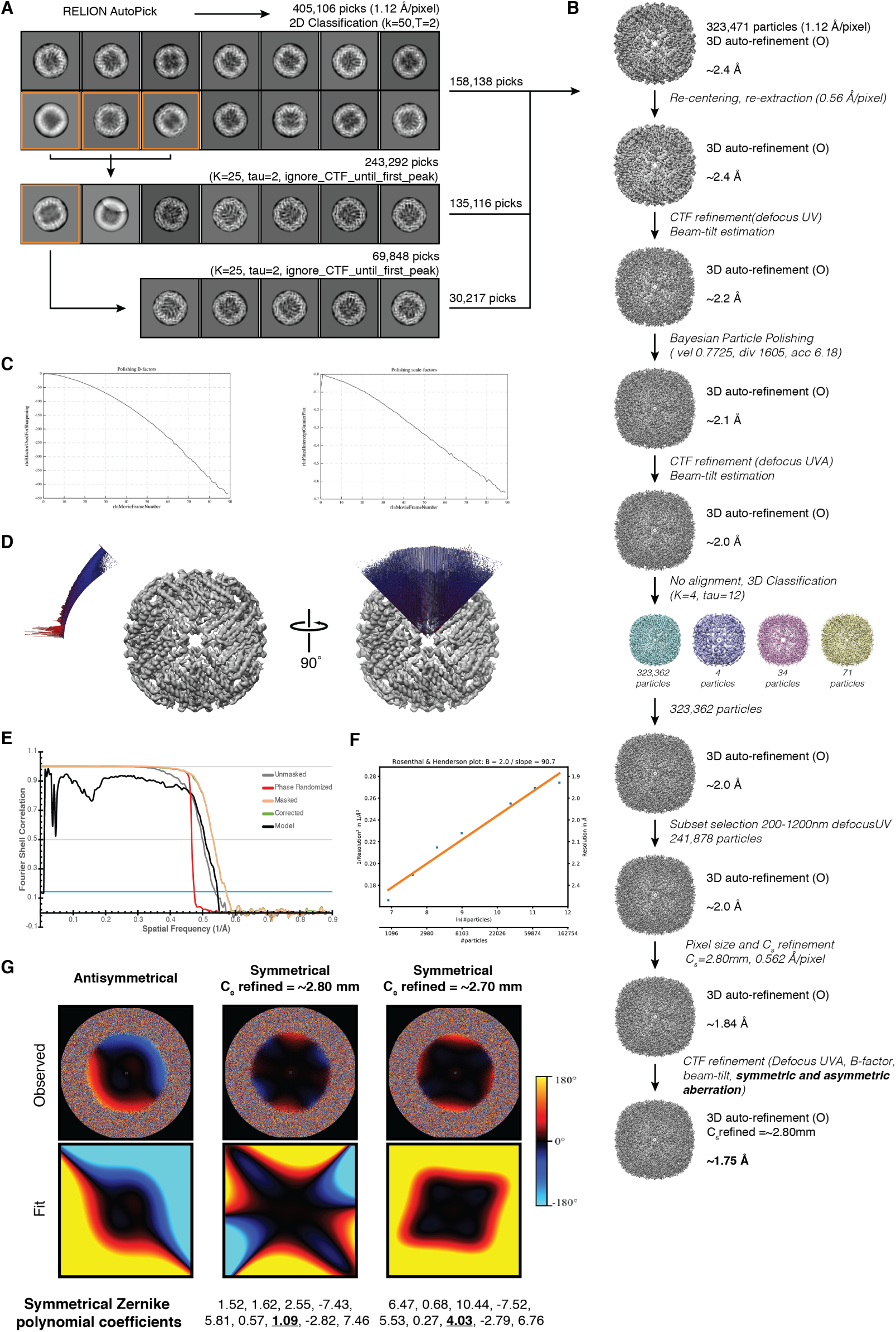
Schematic for apoferritin single-particle cryo-EM data processing. **(A)** ∼400K apoferritin particles were extracted from aligned, non-dose-weighted micrographs, Fourier-binned 2 × 2 (192 pixel box, 1.12 Å/pixel), and subjected to three subsequent rounds of reference-free 2D classification using RELION 3.0^1^. Representative class averages are shown. Particles comprising the “best” classes were saved while the particles contributing to class averages highlighted in orange were subjected to additional rounds of 2D classification. **B)**These particles were combined and subsequently 3D auto-refined to ∼2.4 Å resolution (O symmetry). Particles were re-centered and re-extracted unbinned (384 pixel box, 0.56 Å/pixel), and subjected to 3D auto-refinement followed by CTF refinement (defocus UV, whole micrograph astigmatism), including beam-tilt estimation. After another round of 3D auto-refinement the outputs were used for Bayesian particle polishing to yield “shiny” particles (384 pixel box, 0.56 Å/pixel). Plots of the calculated B-factors and scale factors from Bayesian particle polishing are shown in **(C)**. The outputted “shiny” particles were 3D auto-refined to yield a ∼2.1 Å resolution structure. CTF refinement (defocus UV, whole micrograph astigmatism), including beam-tilt estimation and subsequent 3D auto-refinement (O symmetry) yielded a ∼2.0 Å resolution structure. Particles imaged between 200-1200 nm underfocus were selected (∼240K particles) and 3D auto-refined to ∼2 Å resolution. Changing the spherical aberration (C_s_) value to 2.80 mm and using a refined pixel size of 0.562 Å/pixel followed by CTF refinement (defocus UV, whole micrograph astigmatism) and subsequent 3D auto-refinement yielded a ∼1.8 Å resolution reconstruction. An additional round of CTF refinement including higher-order optical aberration correction and C_s_ refinement yielded a ∼1.7 Å resolution reconstruction. **(D)** Plots showing the Euler angle distribution of the final apoferritin EM density. **(E)** Gold-standard Fourier shell correlation (FSC) curve generated from the independent half maps contributing to the ∼1.7 Å resolution apoferritin reconstruction. FSC curve between the final refined atomic model and apoferritin EM density is also shown. **(F)** Rosenthal and Henderson plot^3^ for particles contributing to the final apoferritin reconstruction. **(G)** Anti-symmetrical and symmetrical aberrations (for refined C_s_=∼2.80 mm and ∼2.70 mm) estimated using RELION 3.1^2^. Per-pixel phase-angle estimates are depicted in the upper plots, and parametric fits using Zernike polynomials are shown in the lower plots. The estimated symmetrical Zernike polynomial coefficients are shown below each panel with the 0,4 term underlined in bold.

**Supplementary Figure 4.**
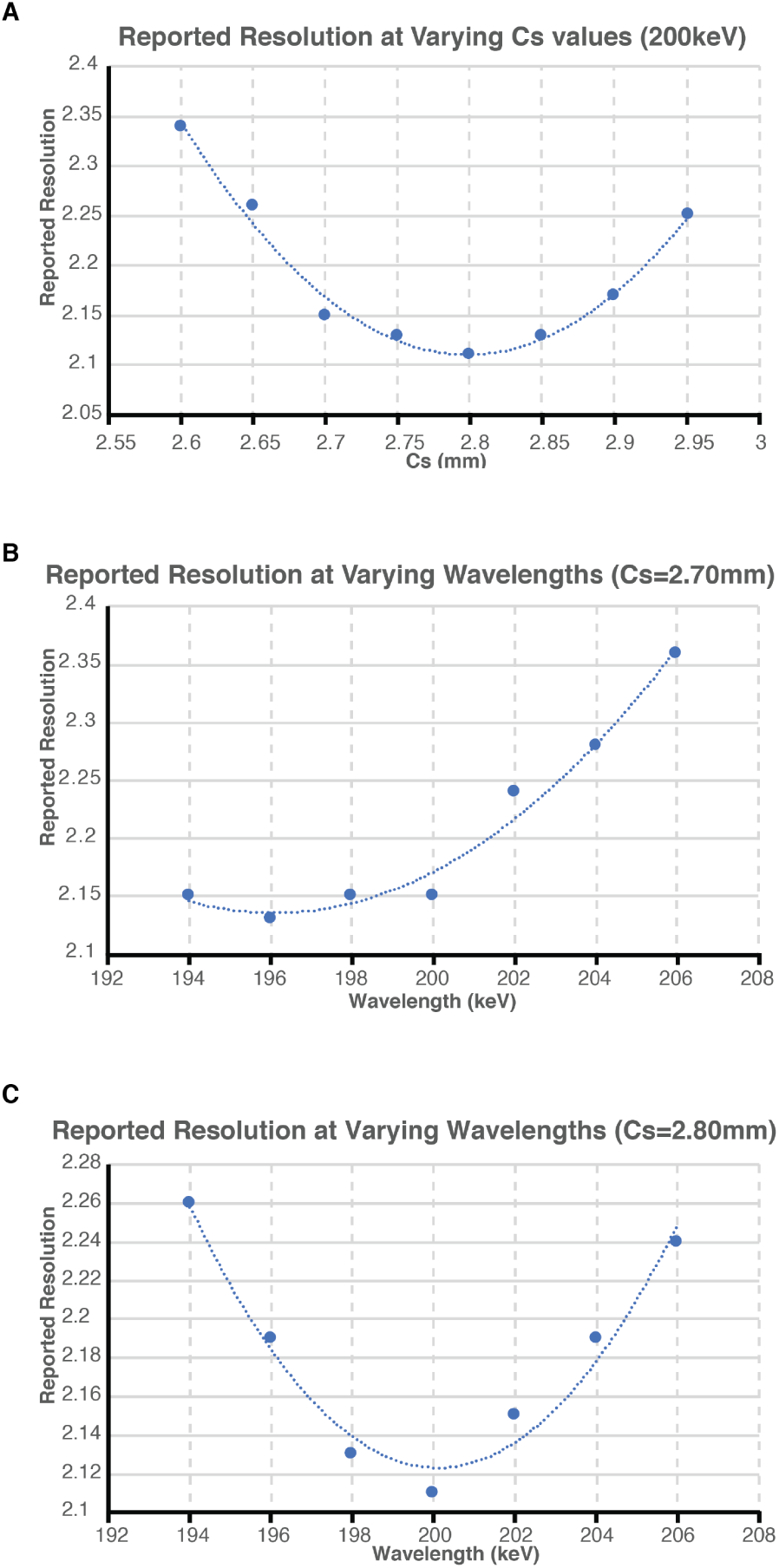
Refinement of spherical aberration and wavelength values. **(A)** FSC-reported resolution values for a subset of the apoferritin data (∼12K particles) in which the C_s_ value was manually changed prior to CTF refinement (defocus UV) and subsequent 3D auto-refinement. Wavelength (200 keV), amplitude contrast (0.1), and astigmatism were kept constant. **(B)** FSC-reported resolution for a subset of the apoferritin data (∼12K particles) where the wavelength was manually changed prior to CTF refinement (defocus UV) and subsequent 3D auto-refinement. The wavelength was varied for subsets using a C_s_=2.80 mm value or **(C)** a C_s_=2.70 mm value. All 3D auto-refinements used the same parameters.

**Supplementary Figure 5.**
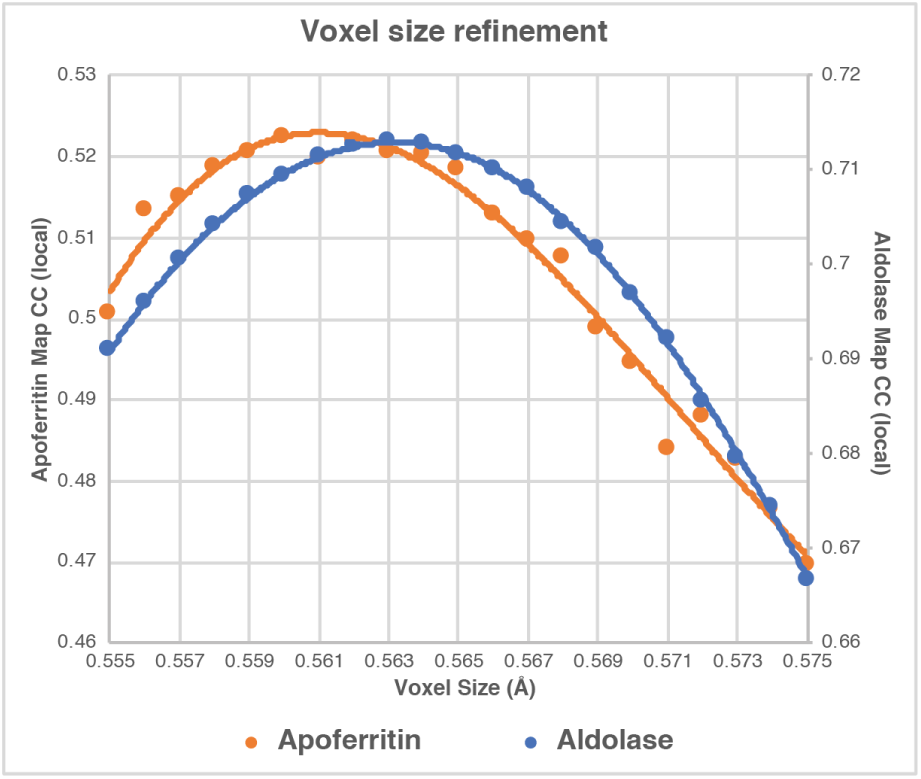
Refinement of map voxel size – aldolase and apoferritin. Measured model-map CC values following Phenix^4^ rigid-body refinement of aldolase (blue dots, PDB ID: 5VY5) or apoferritin (orange dots, PDB ID: 3WNW) asymmetric units (ASUs) into the EM reconstructions of varying voxel size. Polynomial fits for each data set are shown as solid lines.

**Data collection, reconstruction, and model refinement statistics**
|  | Aldolase | Apo ferritin |
| --- | --- | --- |
| **Data collection** |  |  |
| Microscope | Talos Arctica |  |
| Voltage (kV) | 200 |  |
| Nominal magnification | 73,000 |  |
| Exposure navigation | Stage Position |  |
| Cumulative Exposure ($e^- \text{Å}^{-2}$) | 67 | 58 |
| Exposure rate ($e^- \text{pixel}^{-1} \text{sec}^{-1}$) | 1.92 | 2 |
| Exposure per frame ($e^- \text{Å}^{-2}$) | 1.52 | 0.64 |
| Detector | K2 Summit<br>(Counting) | K2 Summit<br>(Super Resolution) |
| Pixel size (Å)* | 0.562 | 0.562 (0.281) |
| Defocus range ($\mu\text{m}$) | -0.3 to -1.8 | |
| Micrographs Used | 3534 | 1759 |
| Total extracted particles (no.) | 1,801,738 | 405,106 |
| Refined particles (no.) | 982,098 | 323,471 |
| **Reconstruction** |  |  |
| Final particles (no.) | 394,294 | 241,878 |
| Symmetry imposed | D2 | O |
| Map sharpening B-factor ($\text{Å}^2$) | -30 | -39 |
| Resolution (global) |  |  |
| FSC 0.5 (unmasked/masked) | 3.06/2.35 | 2.00/1.88 |
| FSC 0.143 (unmasked/masked) | 2.56/2.13 | 1.87/1.75 |
| **Refinement\*\*** |  |  |
| Protein residues | 1372 | 4160 |
| Ligands/waters (atoms) | 328 | >400 |
| Map Correlation Coefficient |  |  |
| Global | 0.77 | 0.63 |
| Local | 0.89 | 0.84 |
| R.m.s. deviations |  |  |
| Bond lengths (Å) | 0.01 | 0.008 ($\pm 0.001$) |
| Bond angles ($^\circ$) | 0.85 ($\pm 0.04$) | 1.23 ($\pm 0.07$) |
| Ramachandran |  |  |
| Outliers (%) | 0.00 | 0.05 ($\pm 0.27$) |
| Allowed (%) | 1.5 ($\pm 0.3$) | 2.3 ($\pm 0.9$) |
| Favored (%) | 98.5 ($\pm 0.3$) | 97.6 ($\pm 1.2$) |
| Poor rotamers (%) | 0.4 ($\pm 0.5$) | 0.3 ($\pm 0.3$) |
| MolProbity score | 1.43 ($\pm 0.10$) | 1.12 ($\pm 0.19$) |
| Clashscore (all atoms) | 6.93 ($\pm 1.21$) | 8.03 ($\pm 1.65$) |
| CaBLAM score<sup>57</sup> | 6.5% ($\pm 0.5\%$) | 1.8% ($\pm 0.5\%$) |
| EMRinger score<sup>58</sup> | 7.15 ($\pm 0.19$) | 8.74 ($\pm 0.10$) |

\*Calibrated pixel size at the detector
\*Values in parentheses correspond to values from top 10 models

